# Nwd1 regulates neuronal differentiation and migration through purinosome formation in the developing cerebral cortex

**DOI:** 10.1101/2019.12.15.877340

**Authors:** Seiya Yamada, Ayaka Sato, Shin-ichi Sakakibara

**Affiliations:** Laboratory for Molecular Neurobiology, Graduate School of Human Sciences, Waseda University, 2-579-15 Mikajima, Tokorozawa, Saitama 359-1192, Japan

**Keywords:** Nwd1, Paics, Fgams, purines, purinosome, neuronal differentiation, neuronal migration, neural stem/progenitor cells, signal transduction ATPases with numerous domains

## Abstract

Engagement of neural stem/progenitor cells (NSPCs) into proper neuronal differentiation requires the spatiotemporally regulated generation of metabolites. Purines are essential building blocks for many signaling molecules. Enzymes that catalyze *de novo* purine synthesis are assembled as a huge multienzyme complex called “purinosome”. However, there is no evidence of the formation or physiological function of the purinosome in the brain. Here, we showed that a signal transduction ATPases with numerous domains (STAND) protein, <u>N</u>ACHT and <u>W</u>D repeat <u>d</u>omain-containing <u>1</u> (Nwd1), interacted with Paics, a purine-synthesizing enzyme, to regulate purinosome assembly in NSPCs. Altered Nwd1 expression affected purinosome formation and induced the mitotic exit and premature differentiation of NSPCs, repressing neuronal migration and periventricular heterotopia. Overexpression/knockdown of Paics or Fgams, other purinosome enzymes, in the developing brain resulted in a phenocopy of Nwd1 defects. These findings indicate that strict regulation of purinosome assembly/disassembly is crucial for maintaining NSPCs and corticogenesis.

## INTRODUCTION

The spatiotemporal differentiation of neural stem/progenitor cells (NSPCs) into immature neurons and neuronal migration are necessary for the proper development of the central nervous system (CNS). The cerebral cortex of embryonic mice contains two distinct types of NSPCs: paired box 6-positive (Pax6^+^) apical progenitor cells (radial glia), located in the ventricular zone (VZ), and T-box brain protein 2-positive (Tbr2^+^) basal progenitor cells (intermediate progenitor cells), which are located the in subventricular zone (SVZ) (Englund et al., 2005). In the neocortex, newborn neurons generated from NSPCs migrate radially toward the cortical plate, accompanied by sequential changes in cell shape. Neurite outgrowth and ensuing polarity formation in immature neurons are also required for cortical layer stratification, and defects in neuronal migration not only cause brain malformation but also various psychiatric disorders, including epilepsy and mental retardation (Hansen et al., 2017; Represa, 2019).

Purines, compounds containing a pyrimidine ring fused with an imidazole ring, are found in all living species and include the nucleobases adenine and guanine (Traut, 1994). Apart from their critical function as the building blocks of DNA (deoxyadenosine and deoxyguanosine) and RNA (adenosine and guanosine), purines work as components of essential biomolecules and as a source of second messenger molecules (cyclic AMP and cyclic GMP), cofactors coenzyme A and nicotinamide adenine dinucleotide (NADH), cellular energy substrate ATP, and GTP which is essential for the signal transduction of a large number of G-proteins. Other purine derivatives contain hypoxanthine, xanthine, and uric acid. Specifically, purines function as neurotransmitters in the brain by acting upon purinergic receptors. Purine metabolites, including ATP and GTP/GDP, are crucial for polarity formation in postmitotic cortical neurons (Raman et al., 2018). During brain development, purinergic signaling is essential for NSPC maintenance and neuronal migration in the neocortical SVZ (Lin et al., 2007; Liu et al., 2008).

In mammalian cells, purine content is regulated by a coordinated balance between the *de novo* and salvage biosynthetic pathways. Although the cellular purine pool is usually supplied by the recycling of degraded bases *via* the salvage pathway, the d*e novo* pathway is upregulated under cellular conditions demanding higher levels of purines, such as tumor growth and cell expansion (Yamaoka et al., 1997). *De novo* purine synthesis comprises a series of 10 enzymatic reactions and is mediated by six evolutionarily conserved enzymes [phosphoribosyl pyrophosphate amidotransferase (PPAT), phosphoribosylglycinamide formyltransferase (GART), formylglycin-amidine ribonucleotide synthase (FGAMS), phosphoribosylaminoimidazole carboxylase phosphoribosylaminoimidazole succinocarboxamide synthetase (PAICS), adenylosuccinate lyase (ADSL), and 5-aminoimidazole-4-carboxamide ribonucleotide formyltransferase inosine monophosphate (IMP) cyclohydrolase (ATIC)], to produce IMP from phosphoribosylpyrophosphate. The enzymes that catalyze *de novo* purine synthesis are assembled near mitochondria and microtubules as a huge multienzyme complex called “purinosome” (An et al., 2010; An et al., 2008; French et al., 2016). Purinosome is a dynamic and functional giant protein complex that emerges during high levels of cellular purine demand in mammalian cultured cells (An et al., 2008). Purinosome formation is linked to cell division (Chan et al., 2015). Furthermore, the dynamic assembly and disassembly of purinosomes *in vivo* might be crucial for the proper development of the human brain. Mutations in *ADSL* and *ATIC* genes cause severe developmental brain defects, such as mental retardation, autistic features, epilepsy, microcephaly, and congenital blindness (Jurecka et al., 2015; Marie et al., 2004). The bifunctional enzyme PAICS, another component of the purinosome, is associated with prostate and breast cancer metastasis and proliferation (Barfeld et al., 2015; Chakravarthi et al., 2018; Meng et al., 2018). PAICS deficiency in humans was recently reported. A missense mutation in *Paics* causes the severe phenotype with multiple malformations, including a small body, short neck, and craniofacial dysmorphism, resulting in early neonatal death (Pelet et al., 2019). To date, however, there is no direct evidence of the localization or physiological function of purinosomes during brain development.

Previously, we identified the NACHT and WD repeat domain-containing protein 1 (*Nwd1*) gene and showed that the Nwd1 protein is expressed in NSPCs and immature neurons in the embryonic mice cerebral cortex (Yamada and Sakakibara, 2018). The Nwd1 protein contains a NACHT domain, which is predicted to have nucleoside-triphosphatase (NTPase) activity, in the central region and a cluster of WD40 repeats at the C terminus. Based on the domain structure, Nwd1 is designated as a novel member of signal transduction ATPases with numerous domains (STAND) protein superfamily and is conserved across species, including zebrafish, mice, rats, monkeys, and humans (Yamada and Sakakibara, 2018). Other members of the STAND protein family often mediate ligand-induced self-oligomerization to form the giant multiprotein complex critical for various important cellular responses; e.g., the apoptotic peptidase activating factor 1 (Apaf1) and nucleotide-binding oligomerization domain-like receptors (NLRs) induce the assembly of large multiprotein complexes, the “apoptosome” and the “inflammasome”, and play central roles in cell death and innate immune responses, respectively (Cai et al., 2017; Dorstyn et al., 2018; Leipe et al., 2004). Although the cellular function of Nwd1 remains unclear, its domain structure is analogous to Apaf1, an essential molecule for apoptosome assembly, which is required for apoptosis initiation (Dorstyn et al., 2018; Yamada and Sakakibara, 2018). This study shows that Nwd1 regulates NSPC proliferation and neuronal migration through the control of purinosome formation during cortical development.

## RESULTS

### Nwd1 Overexpression in vivo Delays the Radial Migration of Immature Neurons and Directs NSPCs to Reside in the VZ

To investigate the role of Nwd1 in the developing cerebral cortex, we overexpressed the *Nwd1* gene *in vivo* using *in utero* electroporation. Full-length Nwd1 or control EGFP was electroporated into NSPCs in the developing dorsal neocortex at E14.5, a stage at which extensive neurogenesis and neuronal migration occurs. Electroporated embryos were harvested and analyzed after 48 h (at E16.5). To visualize the electroporated cells, the EGFP reporter plasmid was co-electroporated with the *Nwd1* plasmid into the same embryos. Figure 1A–1C show that Nwd1 overexpression significantly suppressed neuronal migration from VZ, causing the accumulation of Nwd1-overexpressing cells in VZ/SVZ (control, 26.3%±6.2%, n=6; Nwd1, 73.7%±6.0%, n=6). At E16.5, the majority of cells electroporated with the control EGFP plasmid had migrated and reached the intermediate zone (IZ), whereas Nwd1-overexprssing cells were rarely observed within the IZ (control, 73.6%±6.2%; Nwd1; 26.4%±6.0%) (Figure 1A–1C). Many Nwd1^+^ cells remaining within VZ/SVZ were positive for the neural stem cell marker Nestin (Figure 1D–1H) (control, 29.0%±6.0%, n=4; EGFP-Nwd1, 73.8%±4.8%, n=4), suggesting that they retained their NSPC nature and lined the ventricular wall for at least 2 days, without moving toward the pial surface. After 4 days (at E18.5), EGFP expression was observed in the upper cortical layers and was almost absent in the IZ in controls (Figure 1I). At this time, cells overexpressing Nwd1 remained in the lower layers of the neocortex, including IZ and SVZ (Figure 1I–1K) (layers II–IV: control, 80.5%±1.2%, n=4 vs. Nwd1, 38.4%±6.1%, n=4; layers V–VI: control, 8.3%±1.9% vs. Nwd1, 27.6%±2.7%; IZ: control, 11.3%±1.8% vs. Nwd1, 34.0%±5.7%). Within the lower cortical layers, Nwd1-overexpressing cells exhibited the elongated bipolar morphology of traveling immature neurons. These observations indicated that Nwd1 overexpression directs NSPCs to reside in germinal proliferative zones and delays the radial migration of immature neurons.

**Figure 1.**
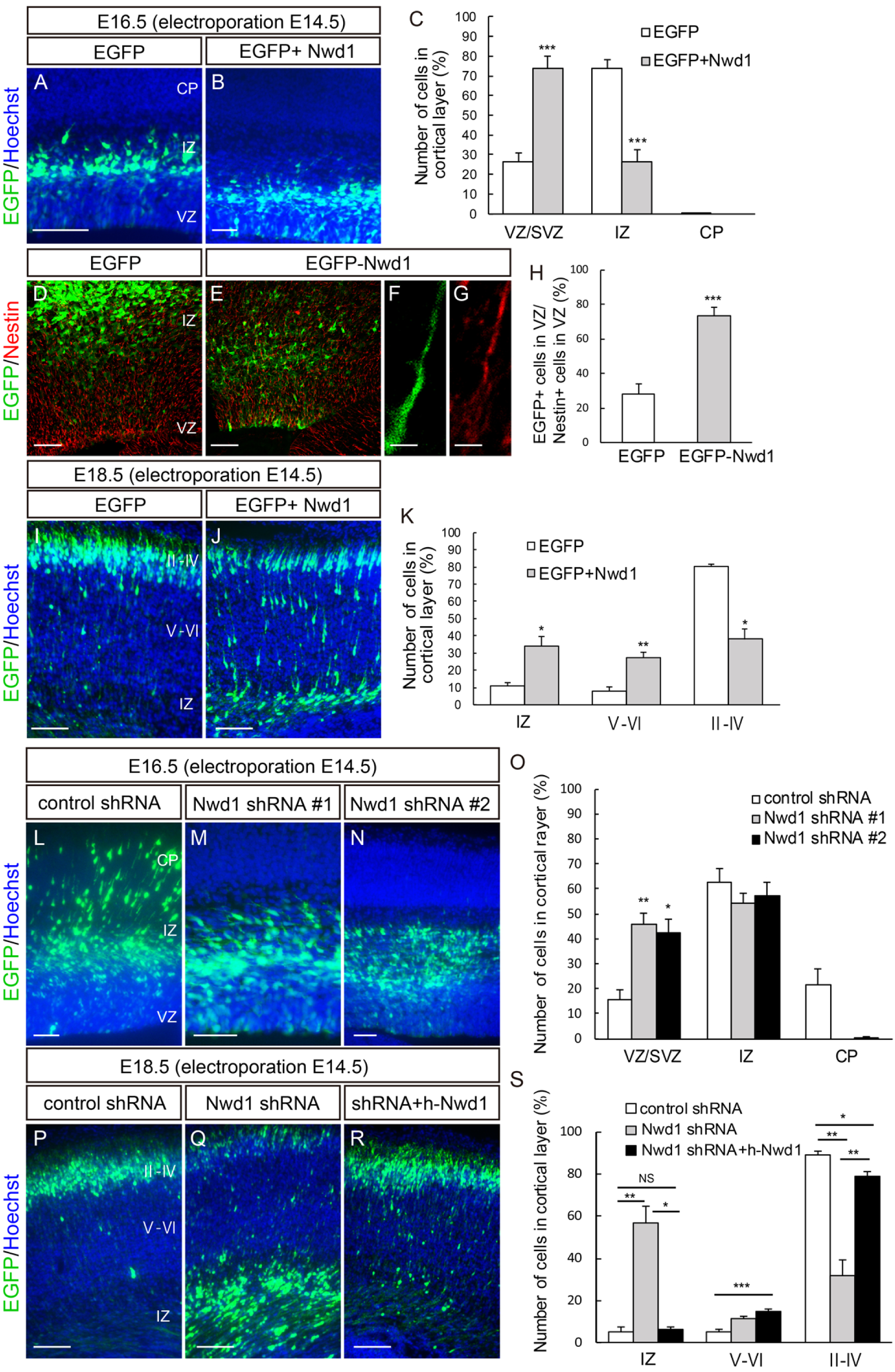
Dysregulation of Nwd1 disturbs the radial migration of neurons and directs NSPCs to reside in the VZ/SVZ. (A–K) Nwd1 overexpression in the embryonic neocortex. (A–C) *In utero* electroporation of control EGFP (A) or Nwd1 together with EGFP (B) was performed on E14.5, and the neocortex was analyzed at E16.5. (C) Distribution of electroporated EGFP^+^ cells in the indicated areas. \*\*\**p*<0.001. (D–H) EGFP-Nwd1 (E) or control EGFP (D) was electroporated at E14.5, and brains were immunostained for Nestin (red) at E16.5. (F, G) Higher magnification of the VZ cells expressing EGFP-Nwd1 (F) and Nestin (G). (H) Ratio of EGFP^+^ or EGFP-Nwd1^+^ cells to the total number of Nestin^+^ cells in the VZ. \*\*\**p*<0.001. (I–K) Nwd1 or control EGFP was electroporated at E14.5, and the brains were collected on E18.5. (K) Distribution of EGFP^+^ cells in the indicated layers. \**p*<0.05, \*\**p*<0.01. (L–S) Nwd1 knockdown represses neuronal migration. An *Nwd1* shRNA (shRNA #1 or shRNA #2) was co-electroporated with EGFP at E14.5 and cortices were analyzed at E16.5 (L–O) or E18.5 (P–S). (O) Distribution of EGFP^+^ cells in the indicated areas at E16.5. \**p*<0.05, \*\**p*<0.01. (R) *Nwd1* shRNA was co-electroporated with the full-length human *NWD1* cDNA at E14.5, and the cortex was analyzed at E18.5. (S) Distribution of EGFP^+^ cells in the indicated areas at E18.5. NS, not significant; \**p*<0.05, \*\**p*<0.01, \*\*\**p*<0.001. All data are presented as means ± m in F and G; 100 μm in I, J, and P–R.

### Nwd1 Knockdown Causes Premature Differentiation of NSPCs and Represses Neuronal Migration

We explored the effect of Nwd1 loss of function on cortical development using small hairpin RNA (shRNA) delivery *in vivo* via *in utero* electroporation. Two different shRNA constructs (shRNA #1 and shRNA #2) targeting the coding region of mouse *Nwd1* significantly reduced Nwd1 protein expression levels (Figure S1A). shRNA specificity was further demonstrated by Nwd1 immunostaining. Endogenous Nwd1 protein expression in cultured NSPCs was silenced by shRNA constructs (Figure S1B–S1E). We co-electroporated one of the shRNA constructs with an EGFP-expression plasmid into the neocortex at E14.5 and harvested embryos after 2 or 4 days. Then, we assessed the distribution of EGFP^+^ cells among the discrete cortical zones. In control embryos, almost all EGFP^+^ cells were found in either IZ or cortical plate (CP), and only a small fraction of cells was observed in VZ/SVZ at E16.5 (Figure 1L). However, *Nwd1* knockdown (KD) resulted in a drastically reduced cell migration into the CP as most cells remained in VZ/SVZ at E16.5 (% of cells in the VZ/SVZ: control, 15.7%±3.7%, n=8; *Nwd1* shRNA #1, 45.9%±4.2%, n=4; *Nwd1* shRNA #2, 42.2%±5.7%, n=5) (Figure 1L–1O). We noticed that *Nwd1* KD cells accumulated in IZ if they were unable to penetrate the boundary between IZ and CP (Figure 1N). This phenotype was more pronounced after a further 2 days of shRNA expression. At E18.5, *Nwd1* KD caused a significant accumulation of cells in IZ (control, 5.4%±2.0%, n=4; *Nwd1* shRNA #1, 56.8%±7.8%, n=5) (Figure 1P–1S). Consequently, fewer cells reached the cortical layers (layers II–IV, control, 89.4%±1.3%, n=4; *Nwd1* shRNA #1, 32.0%±7.1%, n=5; layers V–VI: control, 5.2%±1.0%; *Nwd1* shRNA #1, 11.3%±1.4%) (Figure 1P–1S). These defects were rescued by overexpression of the human NWD1 homolog, which is resistant to targeting by the mouse *Nwd1* shRNA. We co-electroporated *Nwd1* shRNA and the full-length human *NWD1* cDNA into the E14.5 cerebral cortex and performed analysis at E18.5. A large fraction of the electroporated cells reached the upper cortical layers through IZ (Figure 1Q, 1S) (% of cells in layers II–IV, 79.0%±2.4%; layers V–VI, 14.8%±1.3%; IZ, 6.3%±1.6%, n=10), restoring the cellular distribution comparable to that of the non-targeting control (see above). This finding further supported the notion that the loss of function of *Nwd1* causes a severe migratory defect in immature neurons *in vivo*.

We previously reported the substantial expression levels of Nwd1 in VZ/SVZ and immature neurons (Yamada and Sakakibara, 2018). Accordingly, a larger number of cells overexpressing Nwd1 remained within VZ/SVZ (Figure 1). Thus, we examined whether *Nwd1* KD affected the nature of the NSPC pool in VZ/SVZ. At E18.5, i.e., 4 days after shRNA electroporation, double immunostaining revealed that the *Nwd1* KD cells remaining in VZ/SVZ were positive for doublecortin (Dcx) (Figure 2A–2D) and β tubulin III (Figure S2A–S2D), which are markers of newborn immature neurons. Interestingly, *Nwd1* KD drove many VZ cells to ectopically and prematurely express Tbr2, a marker of the SVZ basal progenitor cells (intermediate progenitor cells) (Figure 2E–2H′). Concurrently, we observed a decreased density of Pax6^+^ apical progenitors in the VZ region, where *Nwd1* shRNA was expressed (Figure 2I–2L). Immunostaining for the mitotic marker Ki67 revealed that the proliferation rate of NSPCs was significantly reduced by *Nwd1* KD (control, 27.1%±1.7%, n=6; shRNA #1, 14.4%±2.3%, n=6) (Figure 2M–2O). Based on the early onset of lineage markers and decreased cell division in VZ, we concluded that the loss of function of *Nwd1* induced the premature differentiation of NSPCs. Abnormally produced progenies might follow neuronal differentiation near their place of birth, without proper cell migration.

**Figure 2.**
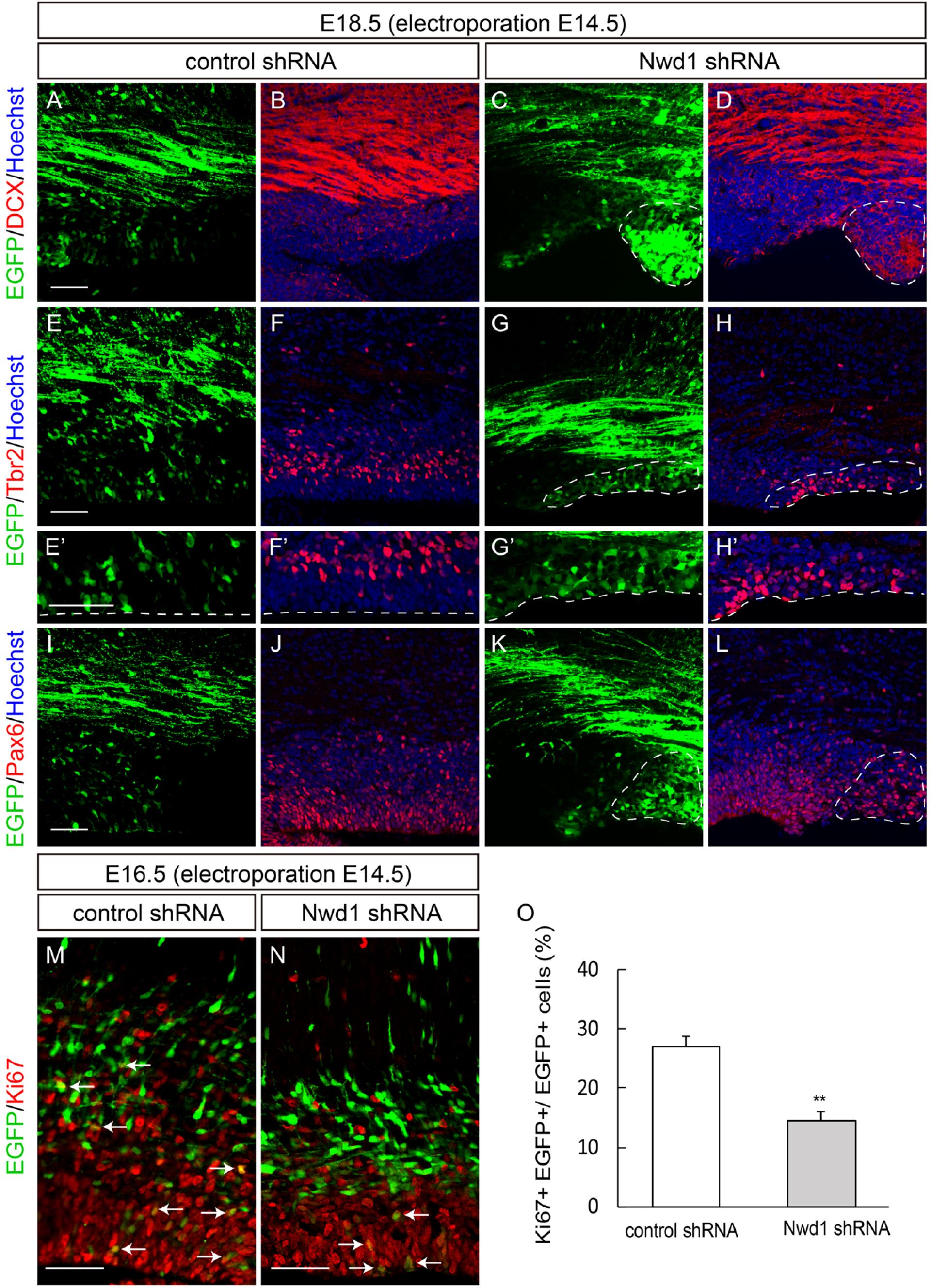
Nwd1 knockdown causes premature differentiation of NSPCs. (A–L) A control or *Nwd1* shRNA was electroporated together with EGFP at E14.5, and the cortices were harvested at E18.5. Confocal images of sections stained with anti-Dcx (A–D), anti-Tbr2 (E–H’), and anti-Pax6 (I–L). The areas surrounded by dashed lines denote the distribution of cells that were electroporated with the *Nwd1* shRNA (green), which remained in ZV/SVZ areas at E18.5. (E’–H’) Higher magnification of the VZ cells shown in (E–H). The VZ surface is outlined by the dashed line. (M–O) An *Nwd1* shRNA was co-electroporated with EGFP at E14.5, and the cortices were stained for the Ki67mitotic marker (red) at E16.5. The arrows indicate EGFP^+^ Ki67^+^ cells. (O) Quantification of EGFP^+^ Ki67^+^ cells in relation to the total number of EGFP^+^ cells. \*\**p*<0.01. Data are presented as means ± SEM. Scale bars, 50 μm.

### Nwd1 Knockdown Causes Periventricular Nodular Heterotopia

We examined postnatal brains after the embryonic KD of *Nwd1*. Embryos electroporated with control shRNA or *Nwd1* shRNA at E14.5 were harvested at postnatal day 7 (P7), when neocortex stratification is almost complete. In a control experiment using a non-targeting shRNA, the electroporated cells were sparsely distributed in the entire cortical region, including the subcortical SVZ (Figure 3A). In contrast, *Nwd1* KD pups frequently developed “periventricular heterotopia,” manifested by ectopic nodules in the lining of the ventricular wall (Figures 3B and S3A–S3C). These heterotopias were characterized by a lower cell density than the neighboring SVZ region and brighter nuclei, evidenced by Hoechst and hematoxylin staining (Figure 3A–3D). Most heterotopia-forming cells had large round cell bodies with few fine processes (Figure 3E, 3H), resembling neurons. Indeed, double immunostaining revealed that they were Dcx^+^ neuron-like cells (Figure 3E–3J). Conversely, these cells never exhibited labeling of the astrocyte marker GFAP (Figure 3K–3P).

**Figure 3.**
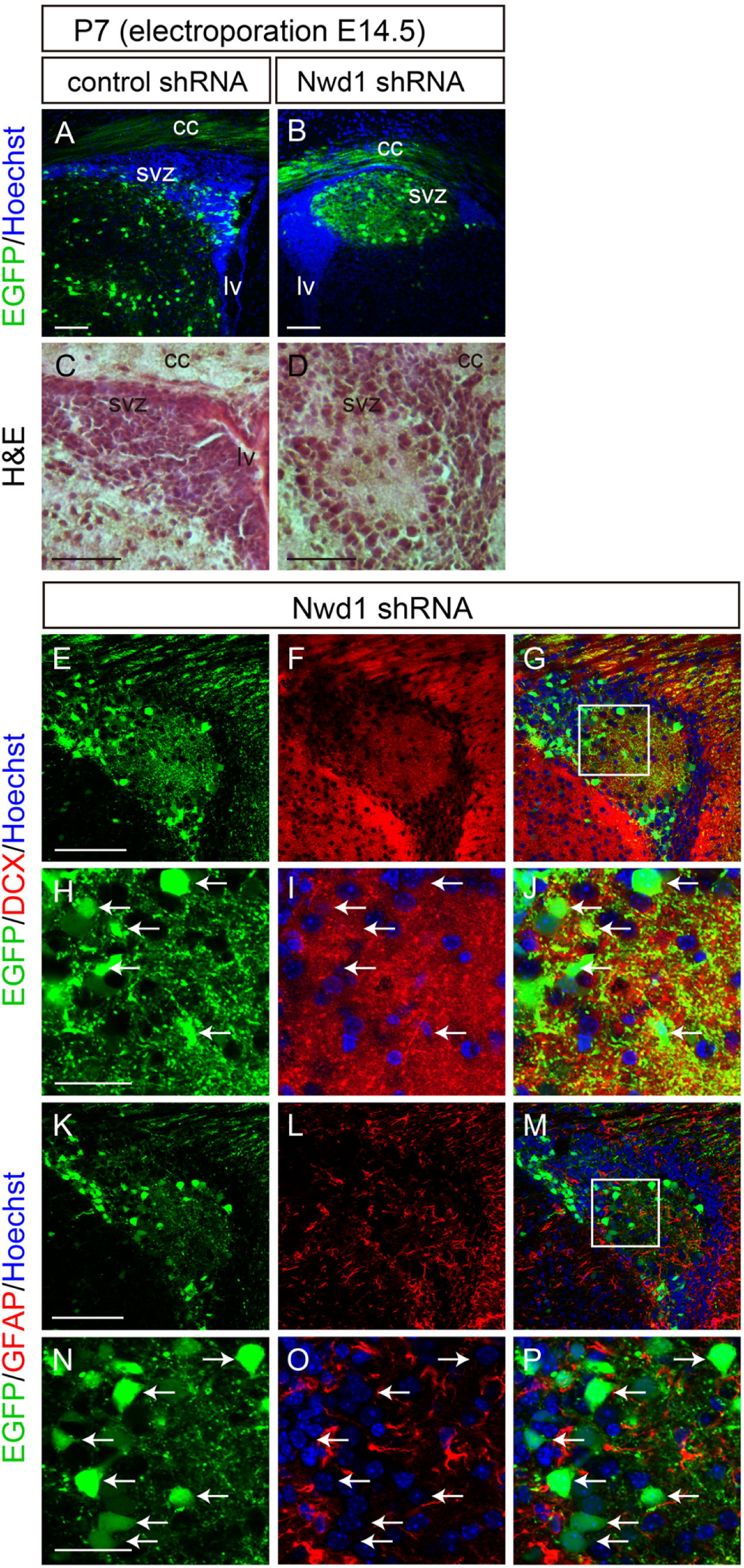
Nwd1 knockdown causes PH. (A–P) A control or *Nwd1* shRNA was delivered into the brain together with EGFP on E14.5, and brains were collected on P7 (n=5). (A, B) Sections through the SVZ area showing the development of periventricular heterotopia (HP) caused by *Nwd1* KD (B). Nuclei are counterstained with Hoechst dye (blue). (C, D) Hematoxylin and Eosin staining of HP. HP regions were immunostained using anti-Dcx (E–J) or anti-GFAP (K–P) antibodies. (H–J) and (N–P) are higher magnifications of the boxed areas shown in (G) and (M), respectively. The arrows indicate the abnormally differentiated neuron-like cells that were Dcx^+^ and GFAP^−^. cc, corpus callosum; lv, lateral ventricle. Scale bars, 100 μm in A–G and K–M; 30 μm in H–J and N–P.

### Expression levels of Nwd1 is Crucial for Neurite Outgrowth and Axon Formation of Cortical Neurons

To understand Nwd1 cellular function in postmitotic differentiating neurons, loss-of-function or gain-of-function experiments were performed using cultured cortical neurons. *Nwd1* shRNA constructs were transferred into dissociated neurons prepared from E16.5 embryos and cultured for 3 days *in vitro* (3 div). In the control, a large fraction (∼75%) of cells extended a single long axon immunostained for the SMI312 neurofilament marker (Figure 4A–4C). At this time, neurons transfected with *Nwd1* shRNA #1 and shRNA #2 exhibited fewer SMI312^+^ axons (Figure 4D–4F), and a notable number of cells lacked axons (Figure 4G). We assessed whether Nwd1 overexpression affected axonal extension in each cortical neuron. Control cells electroporated with EGFP usually had a single axon after culture for 3 div (Figure 4H–4J). In contrast, EGFP-Nwd1 expression inhibited axonal extension (Figure 4K–4M), resulting in apolar-like cells (Figure 4N). These cells occasionally had few short neurites that were devoid of SMI312 immunoreactivity. To visualize all immature neurites extending directly from soma, a plasmid encoding the red fluorescent protein DsRed was co-electroporated into the cortical neurons and the total number of neural processes was counted as neurites. We found that EGFP-Nwd1 overexpression reduced the number of neurites by almost half at 1, 2, and 3 div (Figure 4O–4U) (1 div: control, 4.0±0.2; EGFP-Nwd1, 2.4±0.2; 2 div: control, 4.6±0.2; EGFP-Nwd1, 2.6±0.2; 3 div: control, 5.3±0.1; EGFP-Nwd1, 2.6±0.2). Compared with the control, each neurite appeared thinner and unbranched, suggesting early stage of neurite development. These results indicated that *Nwd1* plays a vital role in the polarity formation in newborn neurons and that tight regulation of its expression level is essential for neurite outgrowth. Recent studies have indicated that dynamic changes in cell shape is closely coupled with the neuronal migration and cortical layer formation (Hirota and Nakajima, 2017). In the developing mammalian neocortex, newborn neurons transiently become multipolar cells with multiple neurites inside SVZ and lower IZ; thereafter, they undergo a change in morphology to a bipolar state before the onset of radial migration to the CP (Maruyama and Okado, 2015); however, its molecular mechanism remains unclear. The defects in neuronal migration caused by manipulating *Nwd1* might reflect a principal function of this gene in the morphological transformation of neurons during neurogenesis.

**Figure 4.**
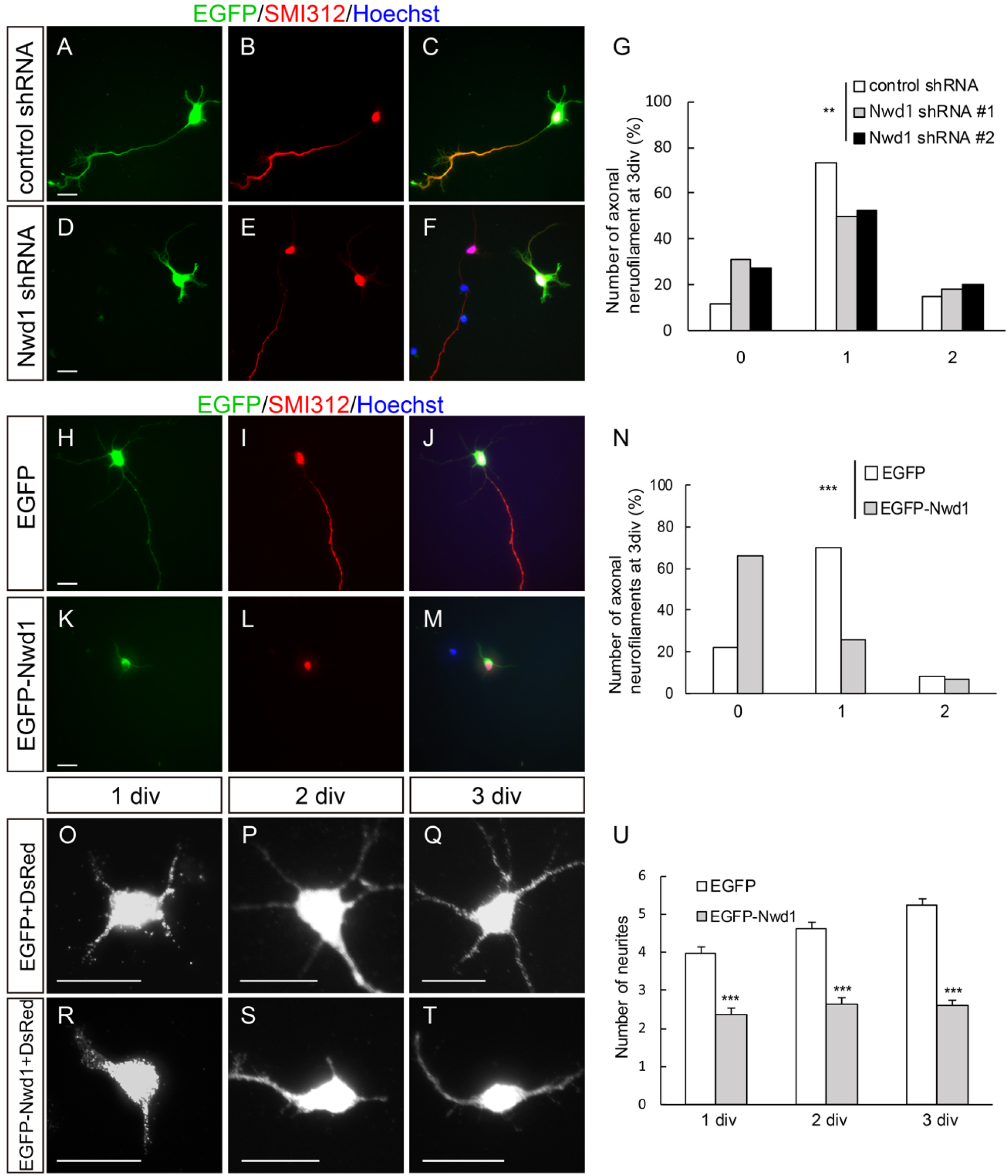
Tightly regulated Nwd1 expression is required for the induction of neuronal identity. (A–G) A non-targeting shRNA or *Nwd1* shRNAs were electroporated together with EGFP into primary cultured cortical neurons. Neurofilaments were stained with a anti-SMI312 antibody (red) at 3 div. (G) Number of SMI312^+^ axons extending from a single neuron. \*\**p*<0.01; control shRNA, n=101; shRNA #1, n=108; shRNA #2, n=99. (H–N) Cortical neurons transfected with EGFP-Nwd1 or control EGFP were stained for SMI312 at 3 div. (N) Number of SMI312^+^ axons extending from a single neuron. \*\*\**p*<0.001; EGFP, n=149; EGFP-Nwd1, n=149. (O–U) EGFP-Nwd1 or control EGFP were electroporated into cortical neurons and cultured for 1 div (O, R), 2 div (P, S), and 3 div (Q, T). To visualize fine immature neurites, a DsRed expression plasmid was co-electroporated into the cells. (U) Number of neurites extending from a single neuron. \*\*\**p*<0.01; EGFP 1 div, n=150; 2 div, n=150; 3 div, n=200; EGFP-Nwd1 1 div, n=150; 2 div, n=150; 3 div, n=200. All data are presented as means ± SEM. Scale bars, 20 μ

### Nwd1 Protein Interacts with Paics

We attempted to understand the molecular mechanism by which Nwd1 regulates cortical development. We used a Y2H screen to identify proteins interacting with Nwd1. Based on its structural similarity to other STAND-family proteins (Leipe et al., 2004), we hypothesized that the N-terminal region of Nwd1 serves as an effector domain by which the protein binds signaling molecule(s) to trigger self-oligomerization mediated by the NACHT domain and WD40 repeats. The N-terminal region of Nwd1 contains a DUF4062 motif, a functionally uncharacterized motif found in bacteria and eukaryotes (Yamada and Sakakibara, 2018). The screening of a mouse brain library using a bait encoding the N-terminal region of Nwd1 led to the isolation of 14 putative Nwd1-binding partners, including Abcd3, Clvs2, Ets1, Paics, Quaking, and Wdr74 (Table S1). Of these binding candidates, Paics was frequently isolated as independent cDNA clones. The interaction between Nwd1 and Paics in yeast was confirmed by the co-transformation of Nwd1 with the rescued Paics plasmid (Figure 5A, 5B). Paics is a bifunctional enzyme that catalyzes *de novo* purine synthesis and is composed of two distinct enzymatic domains: 4-(*N*-succinylcarboxamide)-5-aminoimidazole ribonucleotide synthetase (SAICARs, EC 6.3.2.6) activity in its N-terminal region and 5-aminoimidazole ribonucleotide carboxylase (AIRc, EC 4.1.1.21) activity in its C-terminal region. Since all *Paics* cDNAs identified by Y2H corresponded to the C-terminal region, encompassing AIRc activity domain (Table S1), it is likely that Nwd1 binds to Paics via the this domain (Figure 5C). Nwd1–Paics interaction was further verified by a co-immunoprecipitation (co-IP) experiment using HEK293 cells expressing the Flag-tagged full-length Nwd1 (Flag-Nwd1) and EGFP-tagged full-length Paics (Paics-EGFP). Figure 5D shows that the Paics protein was specifically co-immunoprecipitated with Flag-Nwd1.

**Figure 5.**
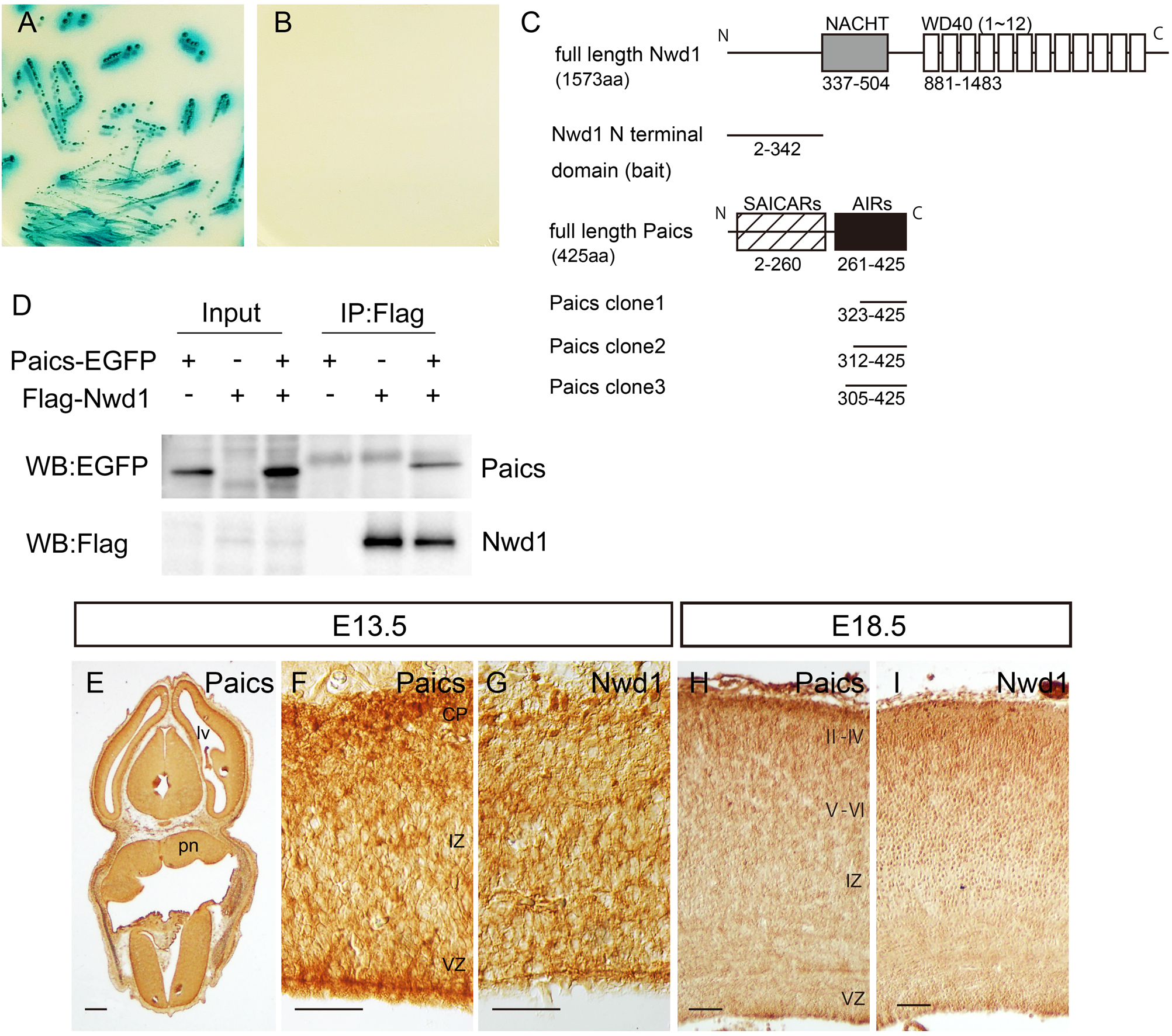
Nwd1 interacts with Paics. (A, B) A yeast transformant with pGBKT7-Nwd1 (bait) and pGADT7-Paics (prey) was streaked on SD agar plates containing quadruple dropout media with Aureobasidin A and X-α-Gal, showing a positive interaction (A, blue colonies). The absence of colonies indicated the negative control (B). (C) Domain structure of the mouse Nwd1 and Paics proteins. The Nwd1 N-terminal region was used as a bait, and the isolated *Paics* cDNAs are indicated. Gray box, NACHT domain; open boxes, WD40 repeats; hatched box, SAICARs activity domain; black box, AIRc activity domain. (D) Co-IP showing the interaction between Nwd1 and Paics. HEK293 cells expressing Flag-Nwd1 and/or Paics-EGFP were subjected to immunoprecipitation with an anti-Flag antibody, followed by immunoblotting with anti-EGFP or anti-Flag antibodies. (E–I) Embryonic brain sections were immunostained with an anti-Paics (E, F, H) or anti-Nwd1 (G, I) antibodies. (E) Transverse section of an E13.5 head. (F–I) Paics (F, H) and Nwd1 (G, I) immunoreactivity in E13.5 or E18.5 cerebral cortex. pn, pons; lv, lateral ventricle. Scale bars, 300 μm in E; 50 μm in F–I.

### Nwd1 and Paics are localized in Purinosomes

We investigated the localization of Paics in the embryonic and postnatal mouse brain. An immunostaining analysis using an anti-Paics antibody showed high levels of expression of Paics in the brain (Figure 5E, 5F). At E13.5, intense immunoreactivity for Paics was detected widely in discrete brain areas, including NSPCs in the VZ/SVZ, in addition to the postmitotic neurons that migrated in the CP of the forebrain, thalamus, midbrain, and hindbrain (Figure 5E, 5F). At E18.5, a lower but detectable Paics expression level was observed in the cerebral cortex (Figure 5H). This spatiotemporal expression pattern of Paics was highly comparable to that of Nwd1 in the developing mouse brain (Figure 5G, 5I) (Yamada and Sakakibara, 2018). Considering the equivalent distribution in the brain and its interaction with Paics, we hypothesized that Nwd1 is involved in the formation of the purinosome. To investigate the localization of Nwd1 in purinosomes, we examined the colocalization of Nwd1 with Paics or Fgams, both of which are used widely as purinosome markers. Because it is technically difficult to detect endogenously formed purinosomes, exogenously introduced markers (such as Fgams-EGFP) are generally used to label intracellular purinosomes (Pedley and Benkovic, 2017). Thus, HeLa cells that expressed Flag-Nwd1 and Paics-EGFP or Fgams-EGFP transiently were cultured in purine-depleted media, to induce the formation of cellular purinosomes (An et al., 2008). Both the Fgams-EGFP and Paics-EGFP proteins exhibited a diffuse cytoplasmic distribution in purine-rich medium (Figure 6A, 6G). In the purine-depleted cells, however, many purinosomes became evident as the cytoplasmic clustering of Fgams-EGFP or Paics-EGFP (Figure 6D, 6J), as described previously (An et al., 2008). We observed the confined distribution and co-clustering of Flag-Nwd1 with Fgams-EGFP^+^ or Paics-EGFP^+^ purinosomes (Figure 6E, 6K).

**Figure 6.**
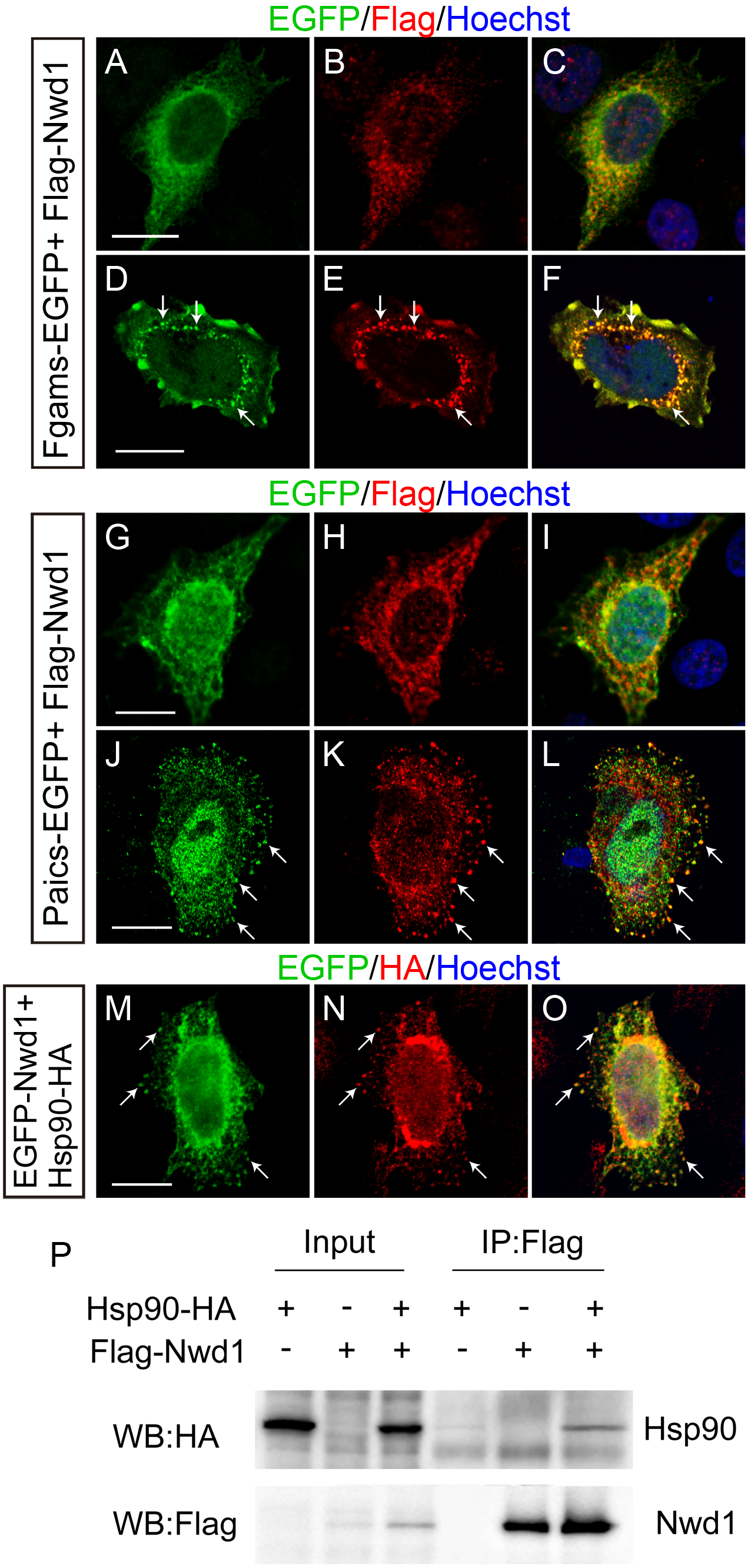
Nwd1 and Paics are localized in purinosomes. (A–L) HeLa cells were transfected with Flag-Nwd1 together with Fgams-EGFP (A–F) or Paics-EGFP (G–L) and cultured in complete medium (A–C, G–I) or in purine-depleted medium (D–F, J–L). Nwd1 was co-clustered with Fgams or Paics as purinosomes in purine-depleted medium (arrows). (M–O) HeLa cells co-transfected with EGFP-Nwd1 and Hsp90-HA were cultured under purine-depleted conditions. The arrows indicate the colocalization of Nwd1 and Hsp90 in purinosomes. (P) Interaction between Nwd1 and Hsp90. HEK293 cells expressing Hsp90-HA and/or Flag-Nwd1 were subjected to immunoprecipitation using an anti-Flag antibody, followed by immunoblotting with an anti-HA or anti-Flag antibody. Scale bars, 15 μm.

A previous study that used co-IP with FGAM followed by a proteomics analysis demonstrated that the heat-shock protein 90 (Hsp90) and Hsp70 colocalize in the purinosome (French et al., 2013). The molecular chaperones Hsp70 and Hsp90 are ubiquitously expressed proteins that have many functions, including assisting protein folding and the stabilization of protein complexes (Makhnevych and Houry, 2012); however, the involvement of these proteins in purinosome formation remains unclear (Pedley and Benkovic, 2017). To validate further the localization of Nwd1 in purinosomes, the distribution of EGFP-Nwd1 and HA-tagged Hsp90 (Hsp90-HA) was assessed in purine-depleted cells; we observed the overlapped localization of Nwd1 in purinosomes (Figure 6O). A co-IP assay of HEK293 cells expressing Flag-Nwd1 and Hsp90-HA demonstrated the interaction between Nwd1 and Hsp90 (Figure 6P). Consistently, Correa et al. showed that NWD1 binds to HSP90 in the human prostate cancer cell line LNCaP (Correa et al., 2014). Taken together, these results strongly indicate that Nwd1 is a novel component of purinosomes.

### Purinosome Assembly is regulated by Nwd1 in NSPCs

To date, there is no evidence of the induction of purinosome assembly in nervous tissues. Therefore, next, we investigated whether NSPCs are capable of forming purinosomes and whether Nwd1 localizes in these structures in NSPCs. NSPCs isolated from the E12.5 cerebral cortex and cultured as a monolayer frequently exhibit fine unipolar or bipolar processes, resulting in a morphology that resembles that of neuroepithelial cells in the embryonic VZ. The expression of Fgams-EGFP distinctly emerged as a granular structure (Figure 7D). Immunostaining using the anti-Paics antibody indicated that a significant proportion of the endogenous Paics protein colocalizes in these clusters (Figure 7E, 7F). The colocalization of Fgams and Paics, which are two enzymes that are essential for *de novo* purine biosynthesis, strongly suggested that these clusters are purinosomes in the NSPCs. Purinosomes were often observed within the cellular processes, in addition to the cell body, under the plasma membrane of NSPCs (Figure 7F). Immunostaining with an anti-Nwd1 antibody revealed the localization of the endogenous Nwd1 protein in Fgams-EGFP^+^ purinosomes in NSPCs (Figure 7G–7L). The purinosome localization of Nwd1 became more evident after the introduction of EGFP-Nwd1 into NSPCs (Figure 7N). These data showed for the first time the presence of purinosomes in NSPCs.

**Figure 7.**
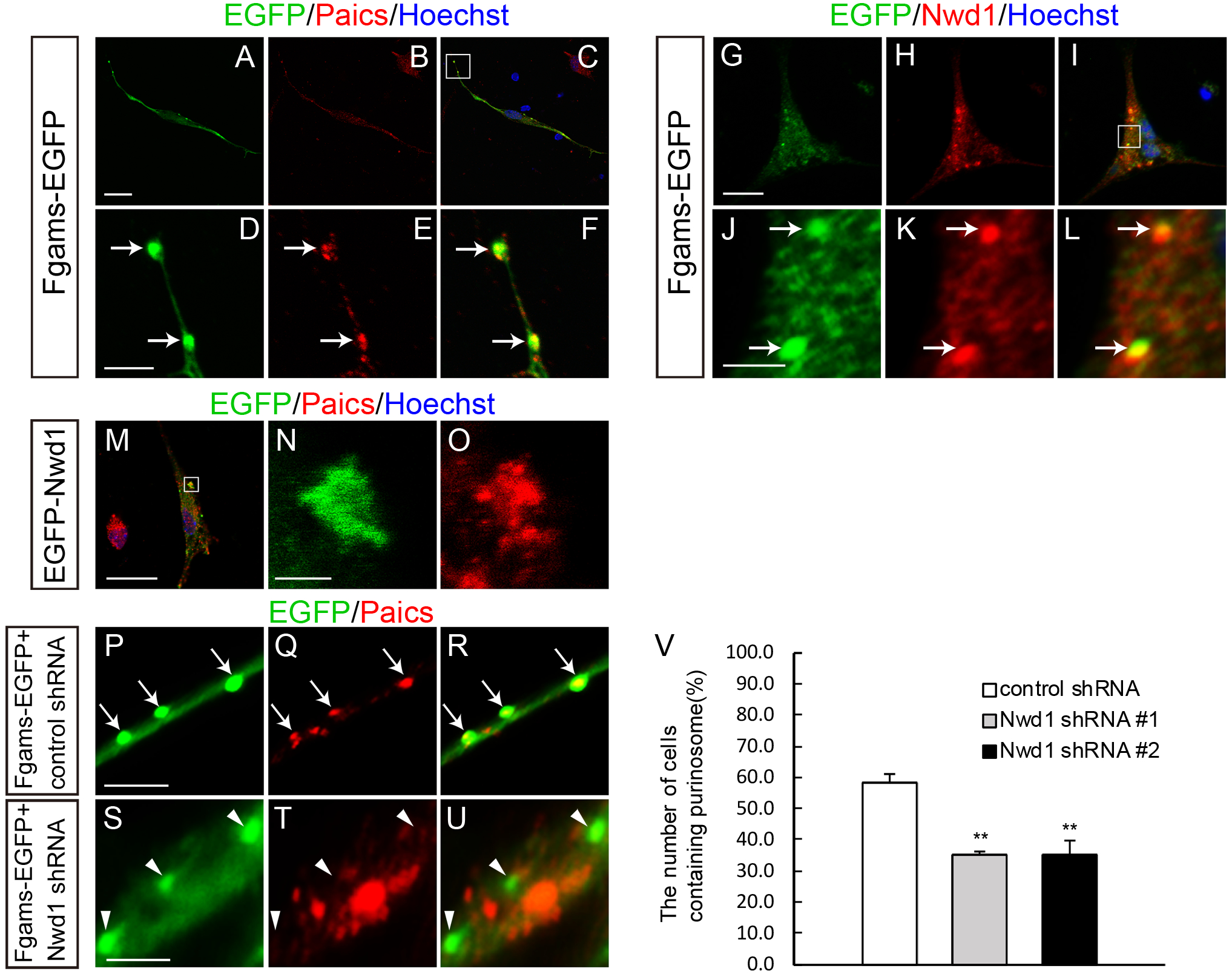
Nwd1 regulates purinosome assembly in NSPCs. (A–C) NSPCs derived from E12.5 telencephalons underwent electroporation with Fgams-EGFP and were immunostained with an anti-Paics antibody at 2 div. (D–F) Higher magnification of the boxed area depicted in (C), showing the clustered signals of Fgams^+^ Paics^+^ purinosomes (arrows) in NSPCs. (G–L) Fgams-EGFP-expressing NSPCs were immunostained with an anti-Nwd1 antibody. (J–L) Higher magnification of the boxed area depicted in (I), demonstrating the localization of endogenous Nwd1 in Fgams-EGFP^+^ purinosomes. (M) EGFP-Nwd1-expressing NSPCs were immunostained with an anti-Paics antibody. (N, O) Higher magnification of the boxed area depicted in (M), showing the colocalization of EGFP-Nwd1 and Paics in purinosomes. (P–V) NSPCs were electroporated with the control shRNA (P–R) or Nwd1 shRNAs (S–U) together with Fgams-EGFP, followed by immunostaining with an anti-Paics antibody at 2 div. The arrows indicate the Fgams-EGFP^+^ Paics^+^ functional purinosomes in NSPCs. The arrow heads indicate the Fgams^+^ Paics^−^ cells. (V) Number of NSPCs containing Fgams-EGFP^+^ Paics^+^ purinosomes. \*\**p*<0.01. Data are presented as means ± SEM. Scale bars, 20 μm in A–C and G–I, M; 4 μm in D–F, J–L, and P–U; 2 μm in N, O.

To examine the role of Nwd1 in purinosome assembly, Nwd1 expression was suppressed by shRNA in NSPCs. *Nwd1* shRNA constructs were electroporated into NSPCs expressing Fgams-EGFP. At 2 div, we counted the number of cells containing Fgams^+^ Paics^+^ purinosomes. As shown in Figure 7V, compared with the non-targeting shRNA, *Nwd1* shRNAs reduced the number of cells containing Fgams^+^ Pacis^+^ purinosomes considerably (control, 58.3% ± 3.0%; shRNA #1, 35.0% ± 1.5%; shRNA #2, 34.7% ± 4.8%). In contrast, the fraction of cells that were labeled with Fgams alone (Fgams^+^ Paics^−^) was increased upon treatment with the shRNAs (Figure 7U). Because a protein complex lacking Paics no longer functions as a purinosome, we concluded that Nwd1 is required for the assembly of the functional purinosome in NSPCs.

### Purinosome Enzymes are Essential for Cortical Development

To clarify the involvement of the purinosome in brain development, we examined the loss-of-function or gain-of-function phenotypes of Paics and Fgams. First, E14.5 embryos were electroporated *in utero* with *Paics* shRNAs, to knockdown the expression of endogenous Paics (Figure S4A). As shown in Figure 8B, *Paics* shRNAs significantly repressed neuronal migration in the neocortex at least until 4 days post-electroporation (E18.5). Compared with the non-targeting control shRNA, *Paics* knockdown resulted in a decreased number of neurons that reached the upper layers II–IV (control shRNA, 73.3% ± 4.8%, n=8; *Paics* shRNA #1, 44.2%±8.2%, n=4; *Paics* shRNA #3, 39.3%±5.8%, n=9) (Figure 8A–8C). Instead, a considerable number of cells accumulated within the IZ and lower cortical layers (V–VI) of embryos expressing *Paics* shRNAs (IZ: control shRNA, 14.4%±3.3%, n=8; *Paics* shRNA #1, 30.7%±7.3%, n=4; *Paics* shRNA #3, 34.0%±6.1%, n=9; layers V–VI: control shRNA, 12.3%±2.4%; *Paics* shRNA #1, 25.1%±1.2%; *Paics* shRNA #3, 26.8%±2.0%; Figure 8C). Notably, 4 days after *Paics* knockdown, a significant number of *Paics* knockdown cells persisted in the VZ/SVZ (Figure 8D). Immunostaining analysis revealed that these VZ/SVZ cells were Dcx^+^, Tbr2^−^, and Pax6^−^ (Figure 8D–8L). Consistently, most of these cells were negative for Ki67 (Figure 8Q), indicating that Paics loss of function induced the mitotic exit and premature differentiation of NSPCs.

**Figure 8.**
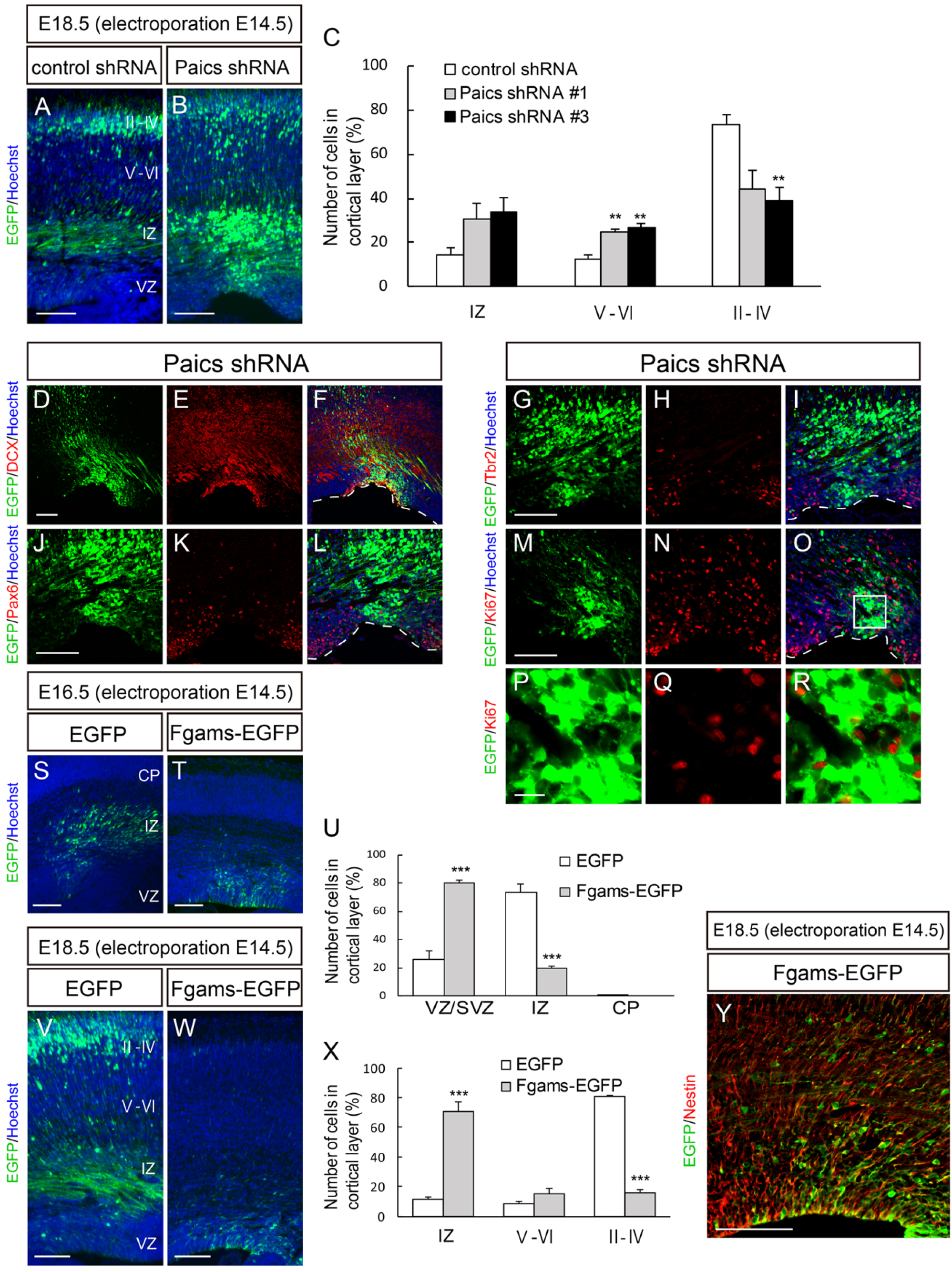
Purinosome components regulate cortical development. (A–R) A control shRNA or Paics shRNA (shRNA #1 or shRNA #3) was delivered into the brain on E14.5, together with EGFP, and the cortices were analyzed at E18.5. (A, B) Distribution of EGFP^+^ cells in the neocortex. (C) Quantification of the distribution of EGFP^+^ cells in the indicated areas. \*\**p*<0.01. (D–R) Brain sections of *Paics* knockdown were immunostained using anti-Dcx (D–F), anti-Pax6 (J–L), anti-Tbr2 (G–I), and anti-Ki67 (M–O) antibodies. The VZ surface is outlined by the dashed line. (P–R) Higher magnification of the boxed area depicted in (O). Many NSPCs electroporated with the *Paics* shRNAs remained in the SVZ/VZ/IZ as postmitotic cells that were Dcx^+^ Pax6^−^ Tbr2^−^. (S–Y) Control EGFP or Fgams-EGFP was delivered into the brain on E14.5 and the cortices were analyzed at E16.5 (S, T) or E18.5 (V, W, Y). (U, X) Distribution of EGFP^+^ cells in the indicated areas at E16.5 (U) or E18.5 (X). \*\*\**p*<0.001. (Y) Fgams-EGFP-overexpressing brain immunostained with an anti-Nestin antibody. Forced expression of Fgams-EGFP directed many NSPCs to remain in the SVZ/VZ as Nestin^+^ cells, and only a few cells reached the border between the SVZ and IZ. All data are presented as means ± SEM. Scale bars, 50 μm in A, B, D, E, and L–N; 5 μm in F and G; 100 μm in I, J, and P–R.

Next, we assessed the effect of Fgams overexpression on neurogenesis and neuronal migration *in vivo*. *Fgams*-EGFP was introduced into NSPCs *in utero* at E14.5. Fgams overexpression significantly suppressed neuronal migration from the VZ, leading to the accumulation of Fgams-overexpressing cells in the VZ/SVZ at E16.5 (control, 26.3%±6.2%, n=6; Fgams, 80.3%±1.9%, n=7) (Figure 8T, 8U). Fgams-overexpressing cells were rarely observed within the IZ (control, 73.6%±6.2%; Fgams, 19.7%±1.9%). At E18.5, most Fgams-EGFP^+^ cells remained in the IZ and SVZ (IZ: control, 11.3%±1.8%, n=4; FGAMS, 70.9%±5.9%, n=8), and fewer cells were found in the cortical neuron layers (Figure 8W, 8X). The Fgams-EGFP^+^ cells that accumulated in the germinal area were Nestin^+^ (Figures 8Y and S5A–S5C), suggesting that they were undifferentiated NSPCs. Taken together, these data provide strong evidence that both Paics and Fgams are essential for neurogenesis and corticogenesis and that the dysregulation of the genes that encode these proteins hinders neuronal migration. Such abnormal properties of neurons caused by the manipulation of Paics and Fgams seemed to be a phenocopy of Nwd1 overexpression/knockdown (Figures 1 and 2). It is likely that the *de novo* biosynthesis of purines, especially the tightly regulated levels of purinosome components, is indispensable for the orchestrated migration and differentiation of neurons that occur during brain development.

## DISCUSSION

### Nwd1 as a Novel Component of Purinosomes

Here, we described for the first time the induction of the formation of Fgams^+^ Paics^+^ purinosomes in NSPCs. We also revealed that Nwd1 interacts with Paics and is localized in purinosomes in NSPCs. Nwd1 functions as a novel component in the assembly of purinosomes. Nevertheless, Nwd1 has no enzymatic activity related to purine biosynthesis, unlike Paics and Fgams. It is possible that Nwd1 participates in the assembly of purinosomes as a member of the STAND family of proteins. The STAND proteins are a newly recognized ATPases associated with diverse cellular activities (AAA) type of ATPases that act as signaling hubs and mediate the energy-dependent remodeling of proteins and the translocation of macromolecules. Generally, *STAND* genes encode multidomain proteins, typically encompassing an N-terminal effector domain, a centrally located NACHT domain which has P-loop ATPase activity and mediates self-oligomerization, and a C-terminal ligand-binding domain (Leipe et al., 2004). The binding of specific ligands onto the C-terminal domain elicits a conformational change in STAND proteins, which is dependent on ATP levels; this results in the formation of the oligomeric ring-shaped superstructures of STAND proteins, which exhibit a central pore (Mermigka et al., 2019). Such superstructures serve as the tightly regulated molecular switch that controls diverse biological processes, including apoptosis and innate immune responses, in which the ring-like superstructures of STAND proteins drive the translocation or remodeling of the substrate proteins (Mermigka et al., 2019).

Among the STAND-family proteins, Nwd1 shares a similar domain structure with Apaf1 (Dorstyn et al., 2018; Leipe et al., 2004; Yamada and Sakakibara, 2018). During apoptosis, the C-terminal WD40 domain of Apaf1 binds to the cytochrome c molecules that leaked from damaged mitochondria. This ligand binding induces the energy-dependent self-oligomerization of Apaf1. Subsequently, the ring-like superstructure of Apaf1 tethers Caspase 9 through the N-terminal CARD domain of Apaf1, to form the macromolecular complex named apoptosome, which triggers the apoptotic caspase cascade (Zou et al., 1999). Similarly, Paics is gathered as a homo-octameric structure in purinosomes (Li et al., 2007), similar to the caspase 9 heptamer in apoptosomes. Based on these observations, we postulated that Nwd1 undergoes an ATP-dependent conformational change upon binding to the ligand(s), via which Paics proteins are recruited and accumulated systematically to form a functional purinosome. Nwd1 may act as a sensor protein that initiates purinosome formation and activates the *de novo* purine biosynthesis pathway during CNS development.

### Purinosome Components Regulate the Maintenance of NSPCs and Neuronal Migration during Cortical Development

We reported previously the strong expression of Nwd1 in NSPCs and immature neurons during the development of the rodent brain (Yamada and Sakakibara, 2018). The present study revealed a similar distribution of Paics and Nwd1 in the developing neocortex. The gain- and loss-of-function of Nwd1, Paics, and Fgams, which were achieved using *in utero* electroporation, demonstrated that these purinosome components are essential for proper cortical development and that their dysregulation leads to a severe delay in the migration of immature neurons. In addition, *in vivo* knockdown of Nwd1 resulted in a decrease in the number of Pax6^+^ apical progenitors, in conjunction with the ectopic emergence of Tbr2^+^ basal progenitor cells in the embryonic VZ. A previous study reported that the forced expression of the Tbr2 transcription factor directs the conversion of radial glia into basal progenitor cells (Sessa et al., 2008). Thus, we assumed that the altered expression level of Nwd1 caused the premature differentiation of NSPCs, suggesting a vital role for this protein in the maintenance of NSPC pools, including CNS stem cells. Consistently, a previous study suggested a possible role for Nwd1 in tumor cells endowed with stem-cell-like properties; *i.e.*, the proliferative and self-renewing properties. The expression of Nwd1is strikingly upregulated by the Sox9 transcription factor in malignant prostate tumor cells (Correa et al., 2014). A gain- and loss-of-function study indicated that Sox9 plays a central role in the specification and maintenance of CNS stem cells that reside in the embryonic VZ and adult SVZ (Scott et al., 2010). As a downstream target of Sox9, Nwd1 may have a function in the maintenance of CNS stem cells. Interestingly, it was also demonstrated that Paics is necessary for the proliferation and invasion of prostate cancer cells and that the silencing of Paics expression abrogates the progression of several types of prostate tumors (Chakravarthi et al., 2018). Taken together with this evidence, our findings imply that the formation of the purinosome machinery is crucial for the maintenance of somatic stem cells and tumor cells, which commonly require a large amount of *de novo* purine production.

In addition, we demonstrated that a tight control of the level of expression of Nwd1 is crucial for neurite extension and the acquisition of cell polarity, and that altered levels of expression of the *Nwd1* gene because migration defects in cortical neurons *in vivo*. Considering that the spatiotemporally controlled outgrowth of neurites is needed for the establishment of neuronal polarity and neuronal migration (Hansen et al., 2017), purinosome formation might be closely linked to the dynamic morphological transformation of migrating neurons that occurs during corticogenesis. Purines affect many aspects of neuronal differentiation. For example, the activation of Rac, which is a small GTP-binding protein, is required for the formation of the leading process in radially migrating neurons in the embryonic cerebral cortex (Konno et al., 2005). A previous *in vitro* quantitative analysis of purines using neuroblastoma cell lines also demonstrated that the content of intracellular purines is increased as the neuronal differentiation proceeds (Gottle et al., 2013), which suggests the association between purine pools and neuronal differentiation. The Nwd1 protein might affect discrete aspects of neural development, including neuronal migration and the maintenance of the NSPC pool, via the regulation of the assembly/disassembly of purinosomes.

### Implication of Nwd1 and Purinosome Components in Neurological Disorders

Downregulation of Nwd1 by shRNA expression in the embryonic cerebral cortex often caused periventricular nodular heterotopia (PH), a cortical malformation that is characterized by the formation of ectopic aggregates of neurons that line the lateral ventricle. These nodules exhibited a rosette-like structure and appeared to be filled with large neuron-like cells that expressed β-tubulin III and Dcx. In humans, PH is associated with intractable epilepsy and intellectual disability (Cossu et al., 2018). Previous studies using a genetic animal model showed that PH is caused by the failure of the radial migration of newborn neurons from the VZ in addition to the abnormal proliferation of NSPCs (Li et al., 2015; Lian and Sheen, 2015); however, the molecular mechanisms underlying the development of PH are not fully understood. Thus, the disturbance of the purine *de novo* synthesis pathway may be associated, at least in part, with the mechanism underlying the pathogenesis of PH.

In addition to PH, purinosome-related genes are responsible for certain neurological disorders. Deficiency of ADSL in humans causes atrophy of distinct regions of the brain, including the cerebral cortex, in addition to hypomyelination and lissencephaly (Jurecka et al., 2015). Patients with ATIC mutation exhibit neurological symptoms, including profound mental retardation and epilepsy accompanied by various dysmorphic features (Marie et al., 2004). A previous study that used cultured fibroblasts from these patients demonstrated that ATIC and ADSL mutations destabilize the assembly of the purinosome to various degrees, and that the ability to form purinosomes is correlated with the severity of the phenotype of individual patients (Baresova et al., 2012). Recently, PAICS deficiency was reported in humans. Patients carrying a homozygous missense mutation in the *PAICS* gene exhibit multiple severe malformations, including a small body and craniofacial dysmorphism, resulting in early neonatal death (Pelet et al., 2019). Although inactivating mutations in the human *NWD1* gene have not been reported to date, it was recently shown that the neuronal expression of NWD1 is upregulated in patients with temporal lobe epilepsy (Yang et al., 2019). Using a mouse model of acute epileptic seizures, it was suggested that Nwd1 regulates the neuronal hyperexcitability of glutamatergic synaptic transmission in the adult brain (Yang et al., 2019). Therefore, Nwd1 might be involved in a novel mechanism of regulation of the synaptic transmission via the formation of purinosomes or other macromolecular complexes.

## ACKNOWLEDGMENTS

This work was supported by JSPS KAKENHI grant number 26430042 (to S.S.) and by Waseda University Grants for Special Research Projects 2014K-6217 and 2015K-249 (to S.S.). We would like to thank Mr. Daisuke Iijima for technical assistance. The authors would like to thank Enago (www.enago.jp) for the English language review.

## AUTHOR CONTRIBUTIONS

All authors had full access to the data in this study and take responsibility for the integrity of the data and accuracy of the analysis. Study concept and design: S. Y., S. S. Acquisition of data: S. Y., A. S., S. S. Analysis and interpretation of data: S. Y., S. S. Drafting of the manuscript: S. Y., S. S. Obtained funding: S. S.

## DECLARATION OF INTERESTS

The authors declare no competing financial interests.

## MATERIALS AND METHODS

### Animals

ICR male mice were purchased from Japan SLC Inc. (Shizuoka, Japan). The date of conception was established by the presence of a vaginal plug and recorded as embryonic day zero (E0). The day of birth was designated as P0. Mice were housed under temperature- and humidity-controlled conditions on a 12/12 h light/dark cycle, with *ad libitum* access to food and water. All protocols were approved by the Committee on the Ethics of Animal Experiments of Waseda University.

### Tissue Preparation

Tissue preparation was performed as described previously (Yamada and Sakakibara, 2018). Embryos at E16.5 and E18.5 were perfused through the cardiac ventricle with 4% paraformaldehyde (PFA) in 0.1 M phosphate buffer (pH 7.4), followed by post-fixation overnight at 4 °C. Fixed embryo brains were cryoprotected in 30% sucrose in phosphate-buffered saline (PBS) overnight at 4 °C and embedded in optimal cutting temperature compound (Sakura Finetek). Frozen sections were cut at a thickness of 14 µm using a cryostat and were collected on MAS coated glass slides (Matsunami Glass).

### Plasmid Vectors

Mouse *Nwd1* cDNAs were subcloned into the pEGFP-C2 vector (Clontech Takara Bio) to express the Nwd1 protein fused with EGFP. The *Nwd1* and *EGFP-Nwd1* cDNAs were subcloned into the pCAGGS vector (a gift from Dr. Jun-ichi Miyazaki, Osaka University, Japan). For the yeast two-hybrid (Y2H) screening, *Nwd1* cDNAs corresponding to the N-terminal portion of the protein (accession number BC082552; 4bp–1026bp) were subcloned into pGBKT7 (Clontech Takara Bio) to express the N-terminal domain of Nwd1 fused with the GAL4 DNA-binding domain. The Flag-tagged human NWD1 expression plasmid was provided by Dr. Correa (Sanford-Burnham Medical Research Institute, Canada) (Correa et al., 2014).

*pFGAMS-EGFP* and *pPAICS-EGFP* (#99108) were gifts from Dr. Stephen Benkovic (Addgene plasmids # 99107 and # 99108, respectively) (An et al., 2008). HSP90-HA was a gift from Dr. William Sessa (Addgene plasmid #22487) (Cardena et al., 1998). *pCAG-DsRED* was a gift from Dr. Connie Cepko (Addgene plasmid # 11151) (Matsuda and Cepko, 2004).

### shRNA Expression Vectors

We purchased a MISSION shRNA vector library encoding the microRNA-adapted shRNA targeting mouse *Nwd1* (Sigma-Aldrich). Among five shRNA clones (TRCN0000257630, TRCN0000247062, TRCN0000257635, TRCN0000257616, and TRCN0000179877), TRCN0000247062 and TRCN0000257635 yielded efficient knockdown of the exogenous Nwd1 and EGFP-Nwd1 expressed in cultured cells; these clones were designated as shRNA #1 and shRNA #2, respectively. The targeting sequences of shRNA #1 and shRNA #2 were: 5′-TACGACTGTGCATGCTCTAAA-3′ and 5′-CAGGTAATCCAAGTTCGATAT-3′, respectively. The two constructs targeted the coding region of the Nwd1 mRNA. We also used MISSION shRNA plasmids for mouse *Paics* (Sigma-Aldrich). Among five clones (TRCN0000076100, TRCN0000076101, TRCN0000076102, TRCN0000076098, and TRCN0000076099), TRCN0000076101, TRCN0000076102, and TRCN0000076098 were designated as shRNA #1, shRNA #2, and shRNA #3, respectively. The targeting sequences for shRNA #1, shRNA #2, and shRNA #3 were 5′-CTGCTCAGATATTTGGGTTAA-3′, 5′-GCTGATGTCATTGATAATGAT-3′, and 5′-GCACCTGCTTTCAAATACTAT-3′, respectively. shRNA #1 and shRNA #2 targeted the coding region, whereas shRNA #3 targeted the 3′ untranslated region (3′-UTR) of the *Paics* mRNA. A non-targeting shRNA (# SHC202) was also purchased from Sigma-Aldrich.

### Primary Antibodies

The following primary antibodies were used: anti-Nwd1 (affinity-purified rabbit polyclonal antibody used previously (Yamada and Sakakibara, 2018), 1:200 for immunostaining, 1:2000 for immunoblotting), anti-Nwd1 (rabbit polyclonal antibody generated by immunizing the recombinant mouse Nwd1 protein; 1:500 for immunostaining, 1:5000 for immunoblotting), anti-Nestin (chicken polyclonal IgY, Aves Labs, NES; 1:4000 for immunostaining), anti-Nestin (rabbit polyclonal, IBL, 18741; 1:250), anti-α-tubulin (rabbit polyclonal, MBL, PM054; 1:2000 for immunoblotting), anti-GFP (chicken polyclonal IgY, Aves Labs, GFP-1010; 1:2000 for immunostaining), anti-GFP (rabbit polyclonal, GeneTex, GTX113617; 1:2000 for immunoblotting), anti-doublecortin (DCX) (goat polyclonal, Santa Cruz, sc-271390; 1:200 for immunostaining), anti-β-tubulin III (chicken polyclonal IgY, AVES Labs, TUJ; 1:1000 for immunostaining), anti-Pax6 (rabbit polyclonal, MBL, PD022; 1:1000 for immunostaining), anti-Tbr2 (chicken polyclonal, Merck Millipore, 633572; 1:1000 for immunostaining), anti-Ki67 (rabbit monoclonal clone SP6, Lab Vision, RM-9106; 1;1000 for immunostaining), anti-Paics (rabbit polyclonal, Proteintech, 12967-1-AP; 1:200 for immunostaining), anti-GFAP (mouse monoclonal clone G-A-5, Sigma-Aldrich, G3893; 1:400 for immunostaining), anti-SMI312 (mouse monoclonal, Biolegend, 837904; 1:1000, for immunostaining), anti-HA (rabbit polyclonal, MBL, 561; 1:200 for immunostaining, 1:2000 for immunoblotting), and anti-DDDDK (Flag) (mouse monoclonal, MBL, FLA-1; 1:10000 for immunostaining and immunoblotting).

### Cell Culture

HEK293T, HeLa, and Neuro2a (N2a) cells were cultured in Dulbecco’s Modified Eagle Medium (DMEM) containing 10% fetal bovine serum (FBS), penicillin/streptomycin, and L-glutamine. In purine-depleted conditions, HeLa cells were cultured in RPMI 1640 medium supplemented with L-glutamine and 5% dialyzed FBS, as described previously (An et al., 2008; French et al., 2013). NSPCs were cultured as described previously (Yamada and Sakakibara, 2018; Yumoto et al., 2013). Briefly, NSPCs were isolated from the E12.5 telencephalon, seeded onto dishes coated with fibronectin and polyethylenimine (PEI) (Sigma-Aldrich), and cultured in Advanced DMEM/F-12 (1:1) (Life Technologies) supplemented with 15 μg/mL insulin (Life Technologies), 25 μg/mL transferrin (Life Technologies), 20 nM progesterone (Sigma-Aldrich), 30 nM sodium selenite (Sigma-Aldrich), 60 nM putrescine (Sigma-Aldrich), 20 ng/mL basic fibroblast growth factor (FGF2) (Merck Millipore), and 10 ng/mL epidermal growth factor (Merck Millipore). For the primary culture of cortical neurons, embryonic cerebral cortices at E16.5 were dissected and mechanically dissociated. After washing with Opti-MEM I (Life Technologies), cells were electroporated, seeded onto poly-D-lysine-coated dishes, and cultured in neurobasal medium containing 2% B27 (Life Technologies) and 1% GlutaMax (Life Technologies) for 1 3 days *in vitro* (div). For immunostaining, cultured cells were fixed with 4% PFA for 20 min at 4°Cand permeabilized in 0.05% Triton X-100 in PBS for 10 min.

### Cell Transfection

Cultured cell lines were transfected with plasmid DNA and PEI MAX (Polysciences) complexes (ratio of DNA to PEI MAX, 1:3, w/w) formed in Opti-MEM I by incubation for 15 min at room temperature. The DNA complexes were added to cell cultures together with Opti-MEM I for 3 h, followed by cultivation with serum containing complete DMEM. Mouse NSPCs were expanded *in vitro* as described above, and primary cortical neurons were electroporated using a NEPA21 Electroporator (Nepagene) according to the manufacturer’s specifications (NSPCs: two pulses of 125 V for 5 ms with an interval of 50 ms; primary cortical neurons, two pulses of 275 V for 0.5 ms with an interval of 50 ms).

### In utero Electroporation

*In utero* electroporation experiments were performed as described previously (Yumoto et al., 2013). Briefly, pregnant mice were anesthetized via intraperitoneal injection of a mixture containing medetomidine, midazolam, and butorphanol, according to a previous protocol (Kawai et al., 2011). A DNA solution (5 μg/μL) in PBS with 0.01% Fast Green dye (Sigma-Aldrich) was injected into the lateral ventricle through the uterus wall, followed by electroporation. The following constructs were electroporated: *pCAG-Nwd1*, *pCAG-EGFP*, *pCAG-EGFP-Nwd1*, *pFgams-EGFP*, *Nwd1* shRNAs, *Paics* shRNAs, and non-targeting shRNA. Electric pulses were generated by NEPA21 (Nepagene) and applied to the cerebral wall using a platinum oval electrode (CUY650P5, Nepagene), with four pulses of 35 V for 50 ms with an interval of 950 ms. An anionic electrode was placed on the lateral cortex, to ensure the incorporation of DNA into the VZ/SVZ. Embryos were perfused at E16.5, E18.5, and P7 with 4% PFA through cardiac perfusion.

### Immunostaining, Western Blotting, and Immunoprecipitation

Immunostaining and western blotting were performed as described previously (Sakakibara et al., 2001; Yamada and Sakakibara, 2018). For immunoprecipitation, cells were washed in ice-cold PBS and lysed in ice-cold lysis buffer containing 50 mM Tris-HCl (pH 7.5), 150 mM NaCl, 2 mM EDTA, 1% NP-40, and protease inhibitors (cOmplete Mini Protease Inhibitor Cocktail, Roche Diagnostics) for 30 min at 4 °C. Lysates were centrifuged at 15,000 rpm for 10 min and the supernatants were incubated with TrueBlot anti-rabbit or anti-mouse IP beads (Rockland) for 1 h (for preclearing). After centrifugation at 15,000 rpm for 10 min, the supernatants were incubated with the primary antibody for 1 h, followed by immunoprecipitation overnight at 4 °C.

Immunocomplexes were washed four times with 0.1% NP40 in PBS, followed by treatment with 2× sample buffer (125 mM Tris-HCl, pH 6.8, 4% SDS, 10% sucrose, and 0.01% bromophenol blue) and immunoblotting.

### Yeast Two-hybrid Screening

To identify Nwd1-binding proteins in the CNS, a Y2H screening was performed using the Matchmaker Gold Yeast Two-Hybrid System (Clontech Takara Bio), according to the manufacturer’s instructions. The sequence encoding the 340 N-terminal amino acid residues of mouse Nwd1 (BC082552; 4–1026 bp) was subcloned in frame into pGBKT7, to express the N-terminal region of Nwd1 (bait) fused with the GAL4 DNA-binding domain. As the prey, we used the normalized mouse brain library (Clontech Takara Bio, Normalized Mate & Plate Library, cat. # 630488), in which each clone was fused with the Gal4 DNA-activating domain (AD). Mated yeast clones were selected using minimal synthetic defined (SD) medium with double dropout (Leu^−^ and Trp^−^) supplement (Clontech Takara Bio) containing Aureobasidin A and X-α-Gal as the blue-colored colonies; this was followed by a second screening using the SD quadruple dropout (Leu^−^, Trp^−^, Ade^−^, and His^−^) selective medium (Clontech Takara Bio). After the elimination of duplicates containing the same AD/library plasmid via yeast-colony PCR, plasmids were rescued from yeast using the Easy Yeast Plasmid Isolation Kit (Clontech Takara Bio). Protein interactions were confirmed by co-transformation of the Nwd1 bait with each candidate prey plasmid into Y2H Gold yeast host cells, followed by sequencing of the cDNA inserts.

### Statistical Analyses

All numerical data are expressed as means ± SEM. In two-group comparisons, Welch’s *t*-test was used to assess the significance of the differences in cell distribution over the cortical layers, the number of neurites, or the number of purinosomes between different groups. In multiple-group comparisons, analysis of variance followed by Welch’s *t*-test was used. The *p* values obtained were corrected for multiple testing using the Holm–Bonferroni correction. The number of axons was compared using the chi-squared test.

## Notes

#### Summary of Updates

REFERENCES list was updated.

