## Supplemental Text for "Nwd1 regulates neuronal differentiation and migration through purinosome formation in the developing cerebral cortex"

**SUPPLEMENTAL INFORMATION**

**Supplemental Table 1.** **Nwd1-binding partners identified by yeast two-hybrid screen**

Each protein entry is shown with the respective NCBI accession number and the identified protein region. The respective protein functions are also listed.

**Supplemental Figure 1. Validation of the *Nwd1* shRNAs**

(A) The non-targeting control or *Nwd1* shRNA (shRNA #1 or shRNA #2) was transfected into HEK293 cells that expressed EGFP-Nwd1 exogenously. The expression level of Nwd1 was evaluated by immunoblotting using an anti-Nwd1 antibody. Immunoblotting using an α-tubulin antibody was performed to ensure the equal loading of cell lysates. (B–E) The control shRNA or *Nwd1* shRNA #1 was transfected into the primary culture of NSPCs, together with EGFP. Cells were immunostained with an anti-Nwd1 antibody at 2 div, which showed that the expression of endogenous Nwd1 (red) was silenced by the *Nwd1* shRNA. Scale bars, 20 μm.

**Supplemental Figure 2. Nwd1 knockdown causes premature differentiation of NSPCs**

The non-targeting control or *Nwd1* shRNA #1 were electroporated into E14.5 brains along with EGFP, and embryos were harvested at E18.5. (A–D) Confocal images of a neocortex stained with an anti-β-tubulin III (red) antibody. The areas surrounded by dahsed lines denote the distribution of cells electroporated with the *Nwd1* shRNA (green) within the VZ. Nuclei are counterstained with Hoechst dye (blue). Scale bars, 50 μm in A–D.

**Supplemental Figure 3. Periventricular heterotopia caused by Nwd1 knockdown**

The *Nwd1* shRNA was electroporated into the neocortex at E14.5 together with EGFP, and brains were collected at P7. Representative confocal image of a coronal section, showing the ectopic formation of periventricular heterotopia composed of densely packed EGFP^+^ cells (A) located under the neocortex (arrow) in the electroporated hemisphere. Nuclei were stained with Hoechst dye (B). Scale bar, 500 μm.

**Supplemental Figure 4. Validation of *Paics* shRNAs**

N2a cells were transfected with three different mouse *Paics* shRNA constructs (shRNA #1, shRNA #2, and shRNA #3) or a non-targeting control shRNA, followed by immunoblotting with an anti-PAICS (A) or anti-α-tubulin (B) antibody.

**Supplemental Figure 5. Forced expression of Fgams-EGFP directs NSPCs to remain in the VZ/SVZ**

Fgams-EGFP was electroporated into the E14.5 neocortex and embryos were analyzed at E16.5. (A) The cerebral cortex was stained with an anti-Nestin antibody (red). (B, C) Magnified view of the boxed area depicted in (A), showing that Fgams-EGFP^+^ cells were persistently located in the VZ/SVZ as Nestin^+^ NSPCs. Scale bars, 100 μm in A; 10 μm in B and C.
