## Supplementary figures and images for "Nwd1 regulates neuronal differentiation and migration through purinosome formation in the developing cerebral cortex"

### Figure S1

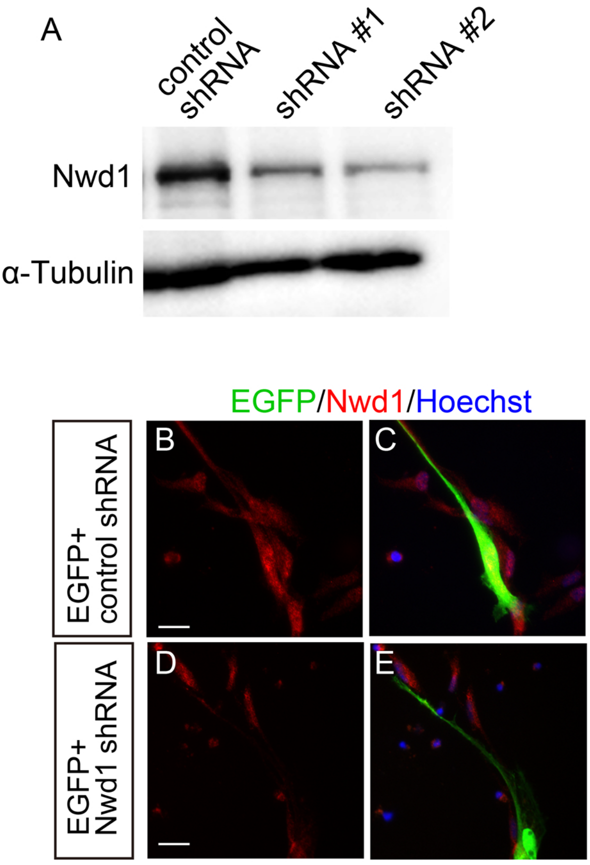

### Figure S2

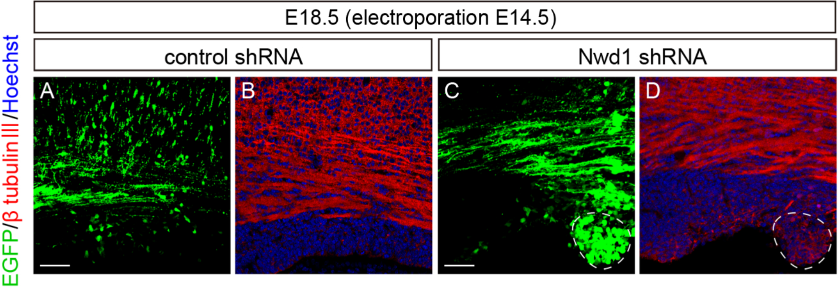

### Figure S3

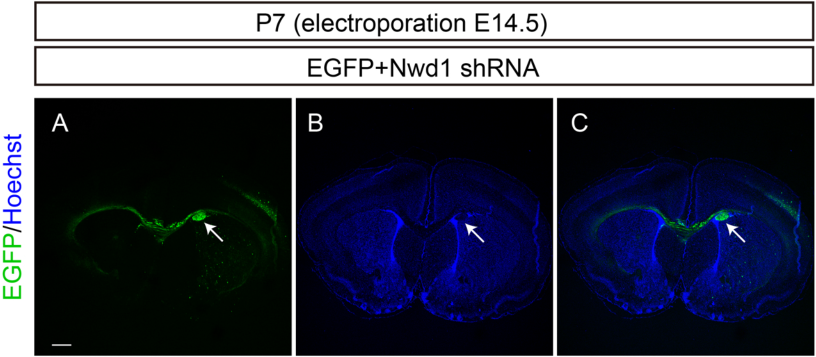

### Figure S4

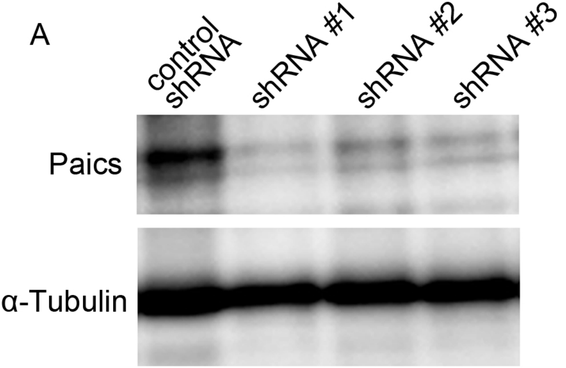

### Figure S5

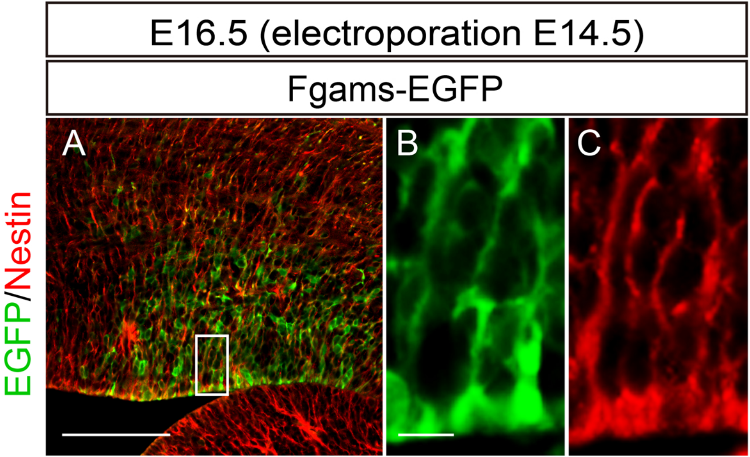
