## Supplementary material for "Nwd1 regulates neuronal differentiation and migration through purinosome formation in the developing cerebral cortex": Table S1

| Gene name | Gene Symbol | NCBI Reference Sequence: protein | Identified protein region (amino acids) | protein function |
|---|---|---|---|---|
| abhydrolase domain containing 3 | Abhd3 | NP_598891.1 | 350–411 aa | unknown, paralog of Abhd1 |
| ATP-binding cassette, sub-family D (ALD), member 3 | Abcd3 | NP_033017.2 | 530–659 aa | peroxisomal import of fatty acids and/or fatty acyl-CoAs |
| chymotrypsin-like elastase family, member 1 | Cela1 | NP_291090.2 | 14–155 aa | protease associated with elastin remodeling |
| clavesin 2 | Clvs2 | NP_001346068.1 | 35–181 aa | recycling of synaptic vesicles |
| Kin17 DNA and RNA binding protein | Kin | NP_079556.1 | 1–188 aa | DNA/RNA binding protein |
| phosphoribosylaminoimidazole carboxylase, phosphoribosylaminoribosylaminoimidazole, succinocarboxamide synthetase | Paics | NP_080215.1 | 323–425 aa, 312–425 aa, 305–425 aa | *De novo* purine synthesis enzymes |
| quaking | Qk | NP_001152988.1 | 130–319 aa | RNA-binding protein |
| serine (or cysteine) peptidase inhibitor, clade E, member 2 | Serpine2 | NP_033281.1 | 307–397 aa | inhibititor for serine proteases |
| serine palmitoyltransferase, small subunit A | Sptssa | NP_598815.2 | 10–71 aa | serine palmitoyltransferase isoenzymes |
| E26 avian leukemia oncogene 1, 5' domain | Ets1 | NP_001359463.1 | 1–111 aa | transcription factor |
| tripeptidyl peptidase II | Tpp2 | NP_033444.1 | 1171–1261 aa | serine exopeptidase |
| WD repeat domain 74 | Wdr74 | NP_598900.1 | 114–378 aa | regulator of exosome complex formation |
